# Evaluation of the inhibitory effects of drugs on the growth of *Babesia* Parasites using real-time PCR method

**DOI:** 10.1101/2020.02.05.935478

**Authors:** Xiao-hu Zhai, Xiao-xiao Feng, Xian Wu, Wei-hua He, Yan-yan Li, Da-wei Yao, Bin Zhang

## Abstract

In order to evaluate the inhibitory effects of drug on the growth of *babesia* parasite, relative quantification real-time PCR method was developed in this study. The *18S rRNA* gene was used as target gene for the 2^−ΔΔCt^ method analysis. Meanwhile, Chicken RNA was added into the parasitized blood for total RNA extraction. The *β-actin* gene of chicken was selected as internal control gene for the 2^−ΔΔCt^ method analysis. Parasitized blood 100 μL, 50 μL, 25μL, 12.5 μL, 6.25 μL was prepared for *B. gibsoni* relative quantification. Regression analysis results revealed that significant linear relationships between the relative quantification value and parasitemia. The *18S rRNA* gene expression was significantly decreased after the treatment of Diminazene aceturate and Artesunate in vitro drug sensitivity test. It suggested that this relative quantification real-time PCR method can be used in evaluating the effects of inhibitory of drug.

## Introduction

*Babesia* is transmitted by ticks to vertebrates and results in serious economic losses in the livestock industry world-wide. Human babesiosis is also worldwide problem and presents a significant health burden in areas where it is endemic [1]. Several antibabesial drugs that have been in use for years such as diminazene aceturate, imidocarb dipropionate, artesunate, atovaquone and atovaquone with azithromycin [2–4]. However, most drugs have proven to be ineffective owing to problems related to toxicity and the development of resistant parasites [4, 5]. Therefore, developing more effective drugs against *babesia* with low toxicity to the hosts has been desired.

At present, there are no recognized standard methods for drug screening or activity evaluation. More recently, in vitro assays have been developed for evaluation of the susceptibilities of *Babesia* to drugs [6–9]. Parasite-infected red blood was diluted with uninfected blood to obtain the blood stock with parasitemia. Then blood with parasitemia were cultured in culture medium containing the indicated concentration of drugs and incubated at 37°C in a humidified multigas water-jacketed incubator. The sensitivity against drugs was evaluated by measuring the rate of parasite growth inhibition. The rate of parasite was calculated by counting the parasitized red blood cells to approximately1000 red blood cells in Giemsia-stained thin blood smears [10]. However, microscopic examination of Giemsia-stained thin blood smears that requires good-quality smears is a time-consuming technique and significant differences in parasitemia estimated by different personnel may be found [11].

The mRNA had been used as a marker of viability, because mRNA is a highly labile molecule with a very short half-life (seconds) [12]. Therefore, mRNA could provide a more closely correlated indication of viability status. Ribosomal RNA (rRNA) has also been selected as an indicator of viability [13] and has been found to be positively correlated with viability under some bacterial-killing regimes [14]. In addition, detection of *16S rRNA* from Chlamydia pneumoniae was demonstrated to provide a more suitable indication of active infection than immunocytochemical detection of specific antigens [15]. In this research, we used *18S rRNA* gene of *B. gibsoni* parasites as molecular target for relative quantification using real-time PCR method.

## Materials and methods

### Chemical reagents

Diminazene aceturate and Artesunate were purchased from Macklin Ltd. (Shanghai, China). Stock solutions of 20 mM in physiological saline were prepared and stored at −20 °C until use.

### Parasite

*B. gibsoni* parasites were isolated from a naturally infected dog in Nanjing, China. This parasite was identified to be *B. gibsoni* according to the 18S rDNA sequences analysis. The blood from sick dog naturally infected with *B. gibsoni* was kept in liquid nitrogen. Then this blood was sub-inoculated into a Beagle. Parasites were first detected in the blood 12 days after inoculation. The blood used in this research was collected form Beagle infected with *B. gibsoni*. Samples were collected in strict accordance with the recommendations in the Guide for the Care and Use of Laboratory Animals of Jiangsu province. The protocol was approved by the Committee on the Ethics of Animal Experiments of JiangSu Agri-Animal Husbandry Vocational College.

### B. gibsoni relative quantification assay

Real-time PCR was used for *B. gibsoni* relative quantification. The 100 μL, 50 μL, 25μL, 12.5 μL, 6.25 μL parasitized blood were prepared. Each blood samples were added 3 μg chicken RNA isolated from spleen tissue and mixed together for RNA extraction. The total RNA of mixture was extracted using RNAiso Plus reagent (Takara, Japan) according to the manufacturer’s instructions. The RNA concentration was measured using a NanoDrop 2000 spectrophotometer (Thermo Fisher Scientific Inc.). Reverse transcription-PCR was conducted using a PrimeScript One-Step RT-PCR kit (Takara, Japan) according to the manufacturer’s instructions. One primer pair (β-acting F 5’-GAGAATTGTG CGTGACATCA-3’ and β-acting R 5’-CCTGAACCTCTCATTGCC A-3’) was used to amplify 157 bp fragment of β-actin gene of chicken [16]. Another primer pair (B.com 339 F 5′-GTCTT-GTAATTGGAATGATGGTGAC-3′ and B.com 339 R 5′-ATGCCCCCAA CCGTTCCTATTA-3′) was used to amplify 339 bp fragment of the *18S rRNA* gene of *Babesia gibsoni* [17]. Real-time PCR was performed with an Applied Biosystems 7500 Real-time PCR System (Applied Biosystems. The cDNA was diluted continuously to make the calibration curves. PCR amplification efficiencies of *B. gibsoni 18S rRNA* gene and Chicken *β-actin* gene were established by means of calibration curves. The expression levels of *18S rRNA* gene, which indicated the quantity of *B. gibsoni,* were analyzed with the 2^−ΔΔCt^ method. Each test was performed in triplicate. The group of 100 μL blood was identified as control.

The quantity of *B. gibsoni* was calculated according to the Eq.1.

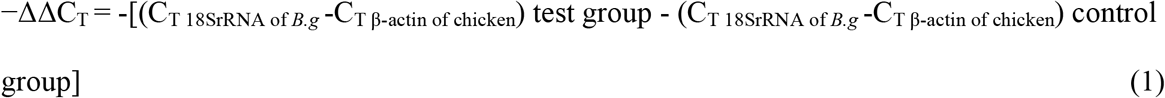

Where C_T 18SrRNA of *B.g*_ was the value of the *B. gibsoni 18S rRNA* gene. C_T β-actin of chicken_ was the value of the Chicken *β-actin* gene.

### Drug sensitivity test

For the in vitro drug sensitivity test, fifty microliters parasitized blood and two hundred microliters of Artesunate (0, 0.005, 0.085 and 0.340 ng/mL, respectively) were distributed per well in 24-well plates in three replicates. These plates were incubated at 37 °C for 24 h. The RNAiso Plus reagent and 3 μg chicken RNA was added into the well for the total RNA extraction. Real-time PCR was used for *B. gibsoni* relative quantification. The group of 0 ng/mL was identified as control. The different concentrations of Diminazene aceturate solution (2500 ng/mL, 250 ng /mL, 25 ng/mL and 2.5 ng/mL, respectively)were used for the in vitro drug sensitivity test as well.

### Statistical analysis

The data in the figure are presented as the arithmetic mean ± standard deviation (SD). The statistical analysis was performed by one-way analysis of variance (ANOVA) using Predictive Analytics Software 18.0. Duncan’s multiple-range test was used, with differences considered to be significant at *P*<0.05.

## Results and discussion

### B. gibsoni relative quantification assay

At present, due to the toxicity and resistance of drugs, the development of new drugs that has a chemotherapeutic effect against babesiosis with high specificity to the parasites and low toxicity to host is urgently needed. Development of a simple in vitro system to evaluate susceptibility of *Babesia* to drugs is very important. Reserve transcription and polymerase chain reaction has proven to be a powerful method to quantify mRNA gene expression. Relative quantification real-time PCR describes the change in expression of the target gene compared to reference group. The method of presenting quantitative real-time PCR data is the comparative CT method, also known as the 2^−ΔΔCt^ method. Using this method, the data are presented as the fold change in gene expression normalized to an endogenous reference gene and relative to the untreated control [18]. For the untreated control sample, ΔΔCt equals zero and 2^0^ equals 1, so that the fold change in gene expression relative to the untreated control equals 1, by definition. Housekeeping genes usually sufficed as internal control genes and it is not affected by experimental treatment. Suitable internal controls for real time quantitative PCR include *GAPDH*, *β-acting*, and *rRNA*.

In this research, *18S rRNA* gene was used quantified the parasites as target gene. It had been reported that the mRNA transcripts of the *B. bovis tubulin* beta chain and small subunit rRNA genes were not affected by the treatment with lower concentration of apicoplast-targeting antibacterials [19]. Due to instability of mRNA, it was sequenced degradation when the parasites were died. Therefore, mRNA could provide a more closely correlated indication of viability status of the parasites.

In this research, *18S rRNA* gene of *B. gibsoni* parasites was selected as target gene rather than internal control genes. Because of degradation of mRNA, there was no gene can be used as internal control genes. In this study, chicken RNA was added into the parasite RNA and the *β-actin* gene of chicken was selected as internal control genes. The same quantification of chicken RNA was added into both treated group and untreated control group to confirm the same quantification expression of *β-actin* gene. The quantity of *B. gibsoni* were calculated according to the equation where −ΔΔC_T_ = −[(C_T 18SrRNA of *B.g*_−C_T β-actin of chicken_) test group − (C_T 18SrRNA of *B.g*_−C_T β-actin of chicken_) control group]. The melting curve of *B. gibsoni 18S rRNA* gene and Chicken *β-actin* gene indicated that the primer used in this study were specific for real-time PCR amplification. PCR amplification efficiencies of *B. gibsoni 18S rRNA* gene and Chicken *β-actin* gene were 97.7% and 96.7%. These results suggested real-time PCR used in this study was reliable. *B. gibsoni* relative quantification was shown in the Fig 1. The magnitude of linearity ranged from 12.5μL to 100 μL. The equation of the parasitized blood volume versus the *B. gibsoni* relative quantification obtained was y=0.01x with a R^2^ of 0.987. Significant linear relationships between the relative quantification and parasitemia were observed.

**Fig 1.**
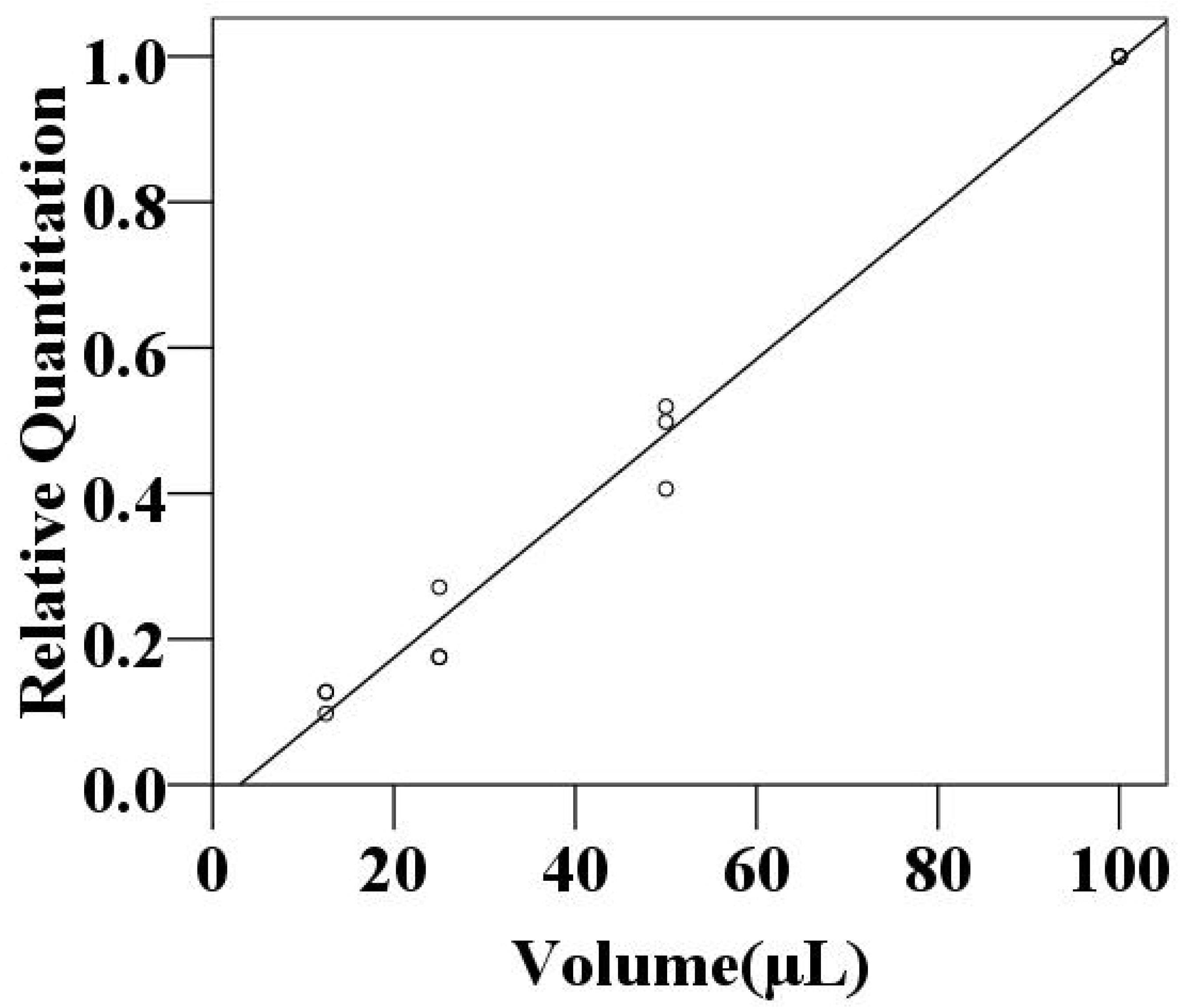
Linearity assessment between *B. gibsoni* relative quantification and parasitized blood volume.

### Drug sensitivity test of Artesunate and Diminazene aceturate

Significant linear relationships between the relative quantification value and parasitemia were observed using the real-time PCR method. In order to evaluate the inhibitory effects of drug, Diminazene aceturate and Artesunate were added into the parasitized blood. It was found that the *18S rRNA* gene expression was decreased relatively to the untreated control. Both Diminazene aceturate and Artesunate can dose-dependently inhibit the growth of *B. gibsoni* in vivo.

The expression levels of *18S rRNA* gene in Artesunate treated group were down-regulated compared to control group (*P*<0.05) (Fig 2). The levels of *18S rRNA* gene were 27% lower at 0.340 ng/mL Artesunate. It was suggested that *B. gibsoni* was significantly suppressed in the presence of 0.005 ng/mL, 0.085 ng/mL and 0.340 ng/mL Artesunate. The higher Artesunate was used, the better the inhibitory effects achieved. Artesunate can dose-dependently inhibit the *B. gibsoni* in vivo.

**Fig 2.**
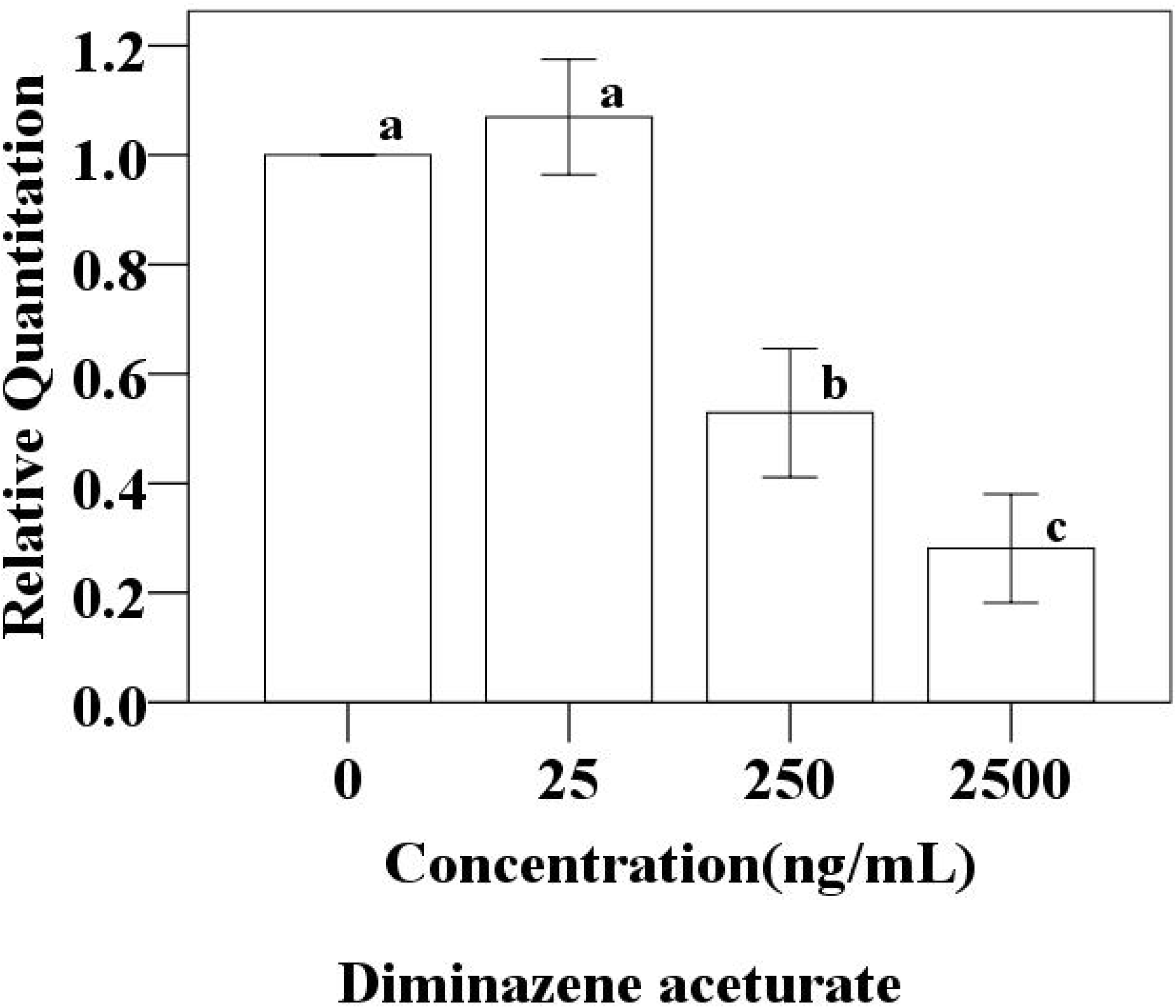
Drug sensitivity test of Artesunate.

The expression levels of *18S rRNA* gene in Diminazene aceturate treated group (250 ng/mL, 2500 ng/mL) were down-regulated compared to control group and 25 ng/mL Diminazene aceturate treated group (*P*<0.05) (Fig 3). The levels of expression were found to be significantly reduce (28%) in the presence of Artesunate2500 ng/mL (*P*<0.05). The lower concentration of Diminazene aceturate (25 ng/mL) had no inhibitory effects.

**Fig 3.**
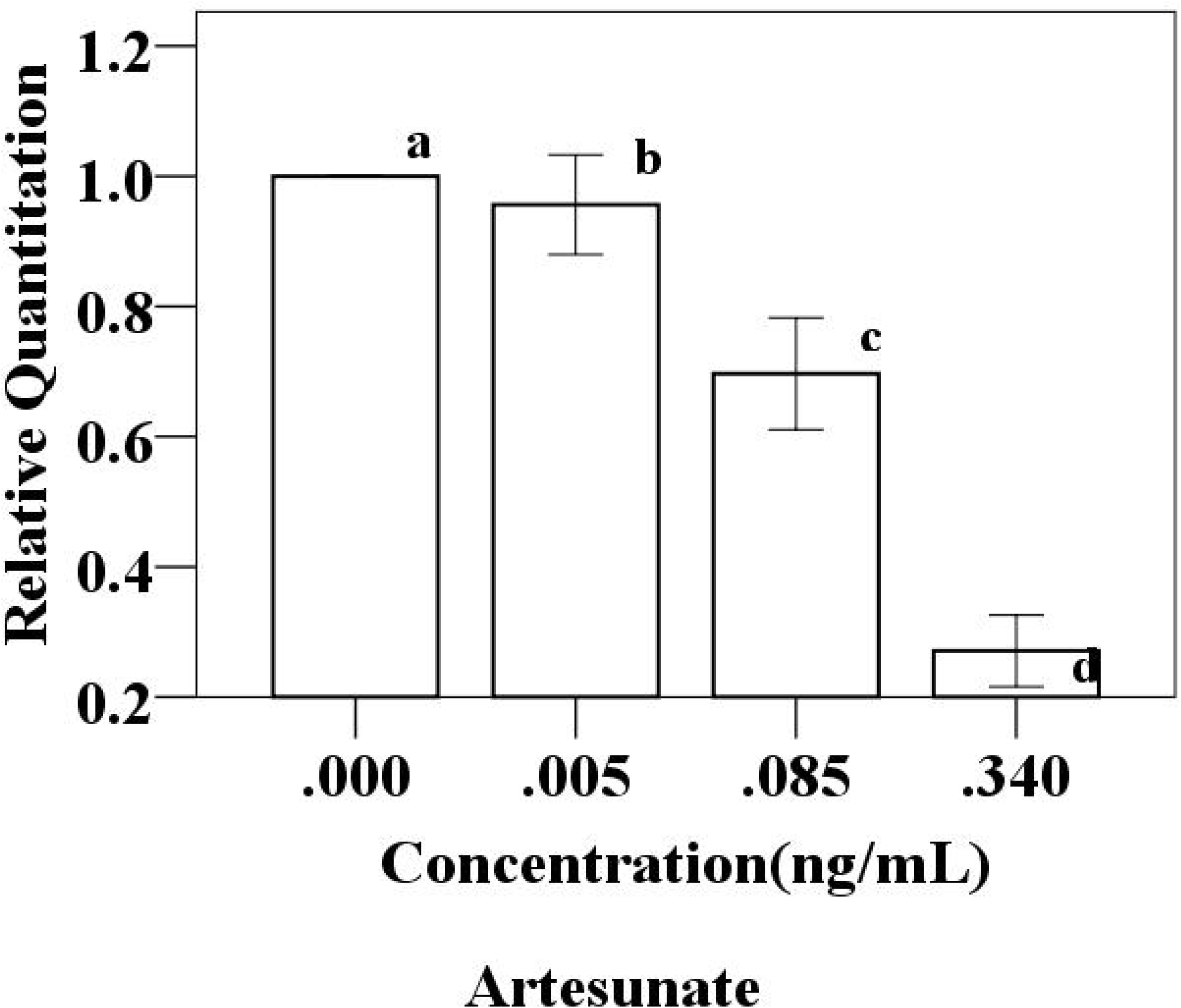
Drug sensitivity test of Diminazene aceturate.

In summary, relative quantification real-time PCR based RNA was developed in this study. This method can determinate live parasites and be used for the evaluation of inhibitory effects of on the growth of *Babesia* Parasites.

## Acknowledgements

We would like to thank Yun Liao and Ze-nan Su for assistance in animal housing.

